# Metabolic reprogramming provides a novel approach to overcome resistance to BH3-mimetics in Malignant Pleural Mesothelioma

**DOI:** 10.1101/2023.03.31.534530

**Authors:** Xiao-Ming Sun, Gareth J Miles, Ian R Powley, Andrew Craxton, Sarah Galavotti, Tatyana Chernova, Alan Dawson, Apostolos Nakas, Anne E Willis, Kelvin Cain, Marion MacFarlane

## Abstract

Malignant pleural mesothelioma (MPM) is an aggressive malignancy linked to asbestos exposure and highly resistant to chemotherapy, potentially due to upregulated expression of the pro-survival proteins, BCL2/BCL-XL/MCL-1. Using clinically-relevant models of MPM we show that patient-derived primary MPM cell lines and *ex-vivo* 3D tumour explants are highly resistant to apoptosis induced by the BCL2/BCL-XL inhibitor, ABT-737. Importantly, we discover that 2-deoxyglucose (2DG), a glycolytic inhibitor, can sensitize MPM cells to ABT-737 and show this correlates with loss of the pro-survival protein, MCL-1. siRNA knockdown of MCL-1 (MCL-1 KD) combined with ABT-737 induced BAX/BAK-dependent, but BIM/PUMA-independent apoptosis, mimicking 2DG/ABT-737 treatment. MCL-1 KD/ABT-737 induced mitochondrial cytochrome *c* release and caspase-independent inhibition of mitochondrial respiration. Moreover, we observed a hitherto unreported caspase-dependent cleavage of glycolytic enzymes and subsequent inhibition of glycolysis. 2DG inhibited ERK/STAT3 activity, decreased MCL-1 mRNA and protein levels, with concurrent activation of AKT, which limited loss of MCL-1 protein. However, co-treatment with a specific AKT inhibitor, AZD5363, and 2DG/ABT-737 potently induced cell death and inhibited clonogenic cell survival, while in MPM 3D tumour explants MCL-1 protein expression decreased significantly following 2DG or 2DG/AZD5363 treatment. Notably, a similar synergy was observed in MPM cell lines and MPM 3D tumour explants using ABT-737 in combination with the recently developed MCL-1 inhibitor, S63845. Importantly, our study provides a mechanistic explanation for the chemoresistance of MPM and highlights how this can be overcome by a combination of metabolic reprogramming and/or simultaneous targeting of MCL-1 and BCL-2/BCL-XL using BH3-mimetics.

## Introduction

Mesothelioma is a malignant tumour with a long latency period, predominately located in the pleural cavity and caused by asbestos inhalation (*1, 2*). Mesothelioma is notoriously refractory to all conventional chemotherapeutic strategies. These regimes provide only marginal benefit and do not significantly increase life expectancy. Therapeutic resistance in tumours can be conferred by mechanisms that activate cell survival signals which promote apoptosis evasion (*3, 4*). Cancer cells also phenotypically exhibit bioenergetic reprogramming *via* the Warburg effect and utilise glucose (glycolysis) more efficiently in an aerobic environment to meet their needs for energy and anabolic intermediates (*5, 6*). This increased uptake and usage of glucose is exploited diagnostically by positron emission tomography scanning to visualise tumours with ^18^F-deoxyglucose (FDG-PET) (*7*). Cell growth signals sustained by oncogenic mutations drive cell-autonomous nutrient uptake and increased proliferative metabolism along with a survival advantage due to increased resistance to apoptotic cell death (*8, 9*). Apoptosis is essential to cellular homeostasis (*10*) and occurs by the extrinsic and intrinsic pathways which transduce death signalling and execute cell death (*11-13*). The extrinsic pathway is activated by diverse death ligands (such as CD95L or TRAIL) binding to their respective death receptors and recruiting intracellular adaptor molecules to form the death inducing signalling complex (DISC) which activates initiator capases-8 or 10 that cleave and activate the executioner caspases-3 and −7 (*14*). However, most environmental stress signals and chemotherapeutic reagents induce cell death *via* the intrinsic pathway, initiated *via* mitochondrial outer membrane permeabilization (MOMP). This results in mitochondrial release of cytochrome *c* along with other apoptotic molecules such as SMAC, OMI or AIF. Cytochrome *c* plays a key role in assembling and activating the Apaf-1/caspase-9 apoptosome complex which initiates caspase activation. MOMP is tightly controlled by BCL-2 family members including pro-survival proteins, such as BCL-2, BCL-XL and MCL-1, which sequester and inactivate pro-apoptotic and pore forming members BAX and/or BAK. A third group of BCL-2 family members, the BH3 only proteins including tBID, BIM, PUMA, NOXA and BAD, either inhibit the pro-survival members or possibly directly activate the pro-apoptotic members (*15*). Thus, over-expression of anti-apoptotic BCL-2 family members is a significant factor in the resistance of cancer cells to chemotherapy. Current strategies have focussed on the development and application of BH3 mimetics to specifically target individual BCL-2 pro-survival proteins, rather than pan-BCL-2 inhibitors which can target multiple family members (*16, 17*). This can result in resistance in cancer cells with higher expression levels of MCL-1 when using BH3 mimetics which are mainly inhibiting BCL-2/BCL-XL. In this respect, we and others have shown that the glycolysis inhibitor, 2-deoxyglucose, (2DG) can downregulate MCL-1, and potentiate apoptosis induced by BCL-2/BCL-XL mimetics and other reagents in certain types of cells (*18-21*).

This combinatorial approach is potentially especially important in MPM cells which express a variety of pro-survival BCL-2 family members (*22*), limiting the use of specific BH3 mimetics. Herein we have used primary MPM cell lines that are highly resistant to ABT-737-induced cell killing and show that 2DG treatment down-regulates MCL-1 and sensitizes primary MPM cell lines to ABT-737-induced cell death. However, 2DG also induces inactivation of ERK/p38 MAPK, potentially independent of RAF/MEK activity and inhibits STAT3 which down-regulates MCL-1 transcription. Conversely, 2DG induced AKT activation which phosphorylates GSK-3β and antagonises MCL-1 degradation and ABT-737-induced cell killing. However, using AZD5363, a specific inhibitor of AKT activity, in combination with 2DG/ABT-737 potentiated clonogenic cell death in both MPM cell lines and in an *ex vivo* 3D MPM explant model. Thus, whilst 2DG sensitizes MPM cells to ABT-737, it also affects other cell signalling pathways which can limit its efficacy. We therefore used MCL-1 KD (siRNA) to specifically recapitulate the direct effects of 2DG on the extrinsic apoptotic pathway and showed that in this scenario (and also with 2DG) ABT-737-induced cell death is BAX/BAK dependent, but BIM/PUMA independent. We also demonstrate that when MCL-1 is down regulated, ABT-737 induces a bioenergetic/metabolic failure (Metabolic Catastrophe; M.C.) by inducing mitochondrial cytochrome *c* release, thereby inhibiting mitochondrial respiration and inducing caspase activation. This induces caspase-dependent cleavage of key glycolytic enzymes and subsequently inhibits glycolysis. We show that combining ABT-737 and S63845 (MCL-1 inhibitor) to simultaneously target BCL-2/BCL-XL and MCL-1 potently induces cell death in both primary MPM cell lines and clinically-relevant *ex vivo* MPM patient-3D tumour explants. Our study provides new insights into drug-resistant mesothelioma but also highlights a potential targetted mechanistic clinical approach to treat this currently incurable disease.

## Materials and Methods

### Derivatisation of cell lines and cell culture

Mesothelioma cell lines MSTO-211H and NCI-H2052 obtained from American Type Culture Collection were cultured in RPMI-1640 supplemented with 10% FBS, L-glutamine (2 mM), penicillin (100 U/ml) and streptomycin (100 μg/ml) and incubated at 37°C under 5% CO_2_ in a humidified atmosphere. MESO-3T, MESO-7T and MESO-8T were derived in our laboratory from mesothelioma tumours after surgical operations with patient consent as previously described (*23*). MPM Cell lines were authenticated by STR profiling commissioned by LGC Standards Cell line authentication team, matching the genomic DNA profile of each patient and not matching any available reference profile (*23*).

### Flow cytometry

After treatment, cells were collected by trypsin (0.5% with EDTA) and suspended in culture medium to allow recovery for 15 min at 37°C. 0.5×10^6^ cells were taken into 0.5 ml of either normal medium (containing 20 nM TMRE) and incubated at 37°C for 10 min for measurement of mitochondrial membrane potential, or Annexin binding buffer (containing Annexin V conjugated with FITC) for phosphatidylserine externalisation assay. Annexin used in this study was sub-cloned, expressed, purified and conjugated in house (*24*). Cells were analysed for induction of apoptosis using a flow cytometer (BD FACSCanto™ II, Beckman Dickenson).

### Western blot analysis

Cells were lysed on ice for 10 min in RIPA buffer containing protease inhibitors (Roche, complete) and phosphatase inhibitors, NaF (5 mM) and Na_3_VO_4_ (2 mM). Cell lysates were clarified by centrifugation at 17,000*g* at 4°C for 10 min. Proteins (20 μg) were separated by SDS-PAGE and transferred to nitrocellulose membranes for western blot analysis as described previously (*23*). Supplementary Table 1 shows a list of Antibodies used in the study.

### siRNA transfection

Transient transfection of siRNA was performed as described by the manufacturer (Life technologies). Briefly, cells were sub-cultured in gelatin-coated plates for at least one day prior to transfection. Cells were transfected by a mixture of siRNA and lipofectamine (RNAiMAX, Invitrogen) in OptiMEM (Gibco) for 48 h prior to the indicated experiments. The siRNAs used in this study are listed in supplemental Table 2.

### CRISPR-Cas9 knockout of BAX and BAK

Lentivirus-based CRISPR-Cas9 genome editing constructs were a kind gift from Prof Stephen Tait (*25*). Viral stocks generated in HEK293T cells were used to infect MESO-8T cells. Single cell colonies were isolated and expanded following trypsinization on cloning discs. Individual cell lines were screened and genomic DNA sequenced to verify indels.

### Cellular metabolism, ATP analysis and cytochrome *c* release assays

Oxidative phosphorylation and glycolysis were simultaneously determined by measuring oxygen consumption rates (OCR) and cellular glycolysis (ECAR, extracellular acidification rate) using Seahorse XF24/96 extracellular flux (XF) analysers (Thermofisher) as previously described (*18, 23*). Briefly, 5 ×10^4^ cells were seeded on gelatin-coated XF24 cell culture microplates at least one day before the experiments. One hour before analysis cells were washed three times with unbuffered and serum-free DMEM containing 1 mM sodium pyruvate, 11 mM glucose and 2 mM glutamine (Glutamax, Gibco) and the final volume adjusted to 675 μL (pH 7.4) before time-dependent analysis. Drugs and inhibitors were loaded into the drug delivery ports and injected as indicated.

Cellular ATP content was measured with Cell-Titer-Glo (Promega) according to manufacturer’s instructions. Resultant chemiluminescence was measured on a 1420 Multilabel Counter (Wallac VICTOR^2^)

Mitochondrial release of cytochrome *c* was detected by selective digitonin permeabilization of the plasma membrane under control conditions as previously described (*11*). Briefly, cells were treated with the indicated drugs and then washed in MitoBuffer (containing HEPES 20 mM, Sucrose 250 mM, MgCl_2_ 5 mM and KCl 10 mM, pH 7.4). Cells were then re-suspended in MitoBuffer containing digitonin, left on ice for 10 min and then centrifuged at 17,000g for 3min at 4°C. Supernatants were collected before resuspending the pellet in MitoBuffer. This mitochondrial fraction along with the supernatants were analysed for cytochrome *c* by SDS-PAGE and western blotting.

### ^35^S incorporation for de novo protein synthesis assay

Cells were treated with 2DG for the indicated times and then incubated in cysteine/ methionine-free DMEM containing ^35^S-methionine (0.37 Mbq/ml, 0.5 ml/treatment) for 20 min then cooled on ice, followed by 3 washes with cold PBS containing unlabelled methionine. Cells were then lysed with Passive Lysis Buffer (400 μl, PLB, Promega) on ice for 5 min, transferred to Eppendorf tubes and centrifuged (1 min) at 17,000*g* at 4°C. The lysates were mixed (1:1 v/v) with 25% TCA and incubated on ice for 30 min. and the precipitated proteins collected on Whatmann Glass Fibre Filters which were air dried for scintillation counting (1414 liquid Scintillation Counter, Wallac)(*26*).

### Quantitative RT PCR

Total RNA was isolated using TRI Reagent (Merck, Gillingham, UK) and. cDNA synthesis performed with random primers and Superscript III (Life Technologies, Paisley, UK). PCR primers were designed as described previously (*27*). Primer sequences are: MCL-1 forward: 5’-ATCTCTCGGTACCTTCGGGAGC-3’, MCL-1 reverse; 5’-GCTGAAAACATGGATCATCACTCG-3’; β2-microglobulin forward: 5’-GCCGTGTGAACCATGTGACT-3’; reverse 5’-GCTTACATGTCTCGATCCCACTT-3’.

Primer concentration were optimized (300– 900 nM) to ensure that the efficiency of the target gene amplification and the endogenous reference amplification were approximately equal. Real-time PCR was performed using SYBR Green PCR Master Mix, primers, and 10 ng of reverse-transcribed cDNA in the PRISM 7500 Sequence Detection System (Applied Biosystems). The thermal-cycler protocol was: stage 1, 50°C/2 min; stage 2, 95°C/10 min; and stage 3, 40 cycles at 95°C/15 s and 60°C/1 min. Each sample was run in triplicate. The CT values for the target amplicon and endogenous control (β2 Microglobulin) were determined for each sample and quantification performed using the comparative CT method (ΔΔCT). Data are presented as the mean ± SEM (n = 3 for each group). Statistical significance was assessed as p < 0.05 using two-tailed student’s *t* test.

### Clonogenicity assay

Cells (5 × 10^3^) were seeded in a 6 well plate and cultured for 14 days with medium changes every 3 days. After treatment, cells were collected by trypsinization and cell number determined using a cell counter (CASY, Scharfe Systems). The cells were washed with PBS and then fixed and stained with 0.75% crystal violet, 50% ethanol, 0.25% NaCl and 1.57% formaldehyde. Excess stain was removed with PBS washing and cell images taken on a LI-COR Odyssey imaging system.

### MPM patient-derived 3D tumour explant model for assessment of drug efficacy

MPM tumour tissue was obtained from surgical operations with patient consent and Research Ethics Committee approval (LREC 08/H0406/226). Explant tissue cubes (2 mm^3^) were cut with skin graft blades on a dental wax plate. Routinely, 8 explants were transferred to a culture plate insert (Millicell®-CM low height, Millipore) and carefully laid on 2 ml culture medium (DMEM containing 2% FBS, 100 U/ml penicillin and 100 μg/ml streptomycin) in 6-well plates. The explants were allowed to recover overnight at 37°C/5% CO_2,_ before incubation for 24 h with fresh culture medium with or without the indicated drugs. The culture media was then replaced with 10% neutral buffered formalin and the explants left at room temperature for ≥ 8 h. The explants were transferred to cassettes and fixed in 70% ethanol and processed to paraffin wax blocks using standard histological procedures. Adjacent tissue sections were de-paraffinized/stained with respective antibodies to cytokeratin (tumour marker), cleaved-PARP (caspase activity) and MCL-1 using standard immunohistochemistry protocols. Immunohistochemistry images were scanned and acquired with a Hamamatsu slide scanner (Fig. 5A & 5D) and then analysed with a Visiopharm Workstation.

## Results

### Inhibition of glycolysis down regulates MCL-1 and sensitises primary patient derived MPM cells to ABT-737-induced apoptosis

MPM is incurable with platinum-based chemotherapy, which remains the preferred treatment (*28*) and therefore to develop new mechanistic based treatments we explored the possible use of BCL-2/BCL-XL inhibitory BH3 mimetics. Initially, we assessed the sensitivity of three newly patient derived primary MPM cell lines (MESO-3T,-7T and −8T) (*23*) and two routinely used commercially available lines (MSTO-211H and NCI-H2052) to ABT-737, a first generation BH3 mimetic (*16*). Most cell lines tested, apart from MESO-3T, were insensitive to high concentrations (≥∼20 μM/6 h) of ABT-737-induced apoptosis (Fig. 1A). Western blot analysis revealed that 8 primary MPM and 6 commercial cell lines predominantly expressed BCL-XL and MCL-1 (Fig. S1A), although interestingly, MESO-3T, −12T and −50T and MSTO-211H also expressed BCL-2 (Fig. S1A, lanes 1, 4, 6 & 10). MESO-7T and −8T had very similar levels of MCL-1 which is not inhibited by ABT-737 (*29*), while MESO-3T has a relatively lower ratio of MCL-1 to Bcl-2, which perhaps explains why these cells are sensitive to ABT-737. Previously we showed that pre-treatment of Z138 cells (mantle cell lymphoma cell line) with 2DG decreased MCL-1 expression and potentiated ABT-737 killing (*18*). As other studies have shown similar results with other cell types (*19*) we reasoned that 2DG could also potentiate ABT-737-induced apoptosis in resistant MPM cells. Significantly, 2DG pre-treatment alone did not induce cell death, but markedly potentiated ABT-737-induced apoptosis, in all resistant primary MPM cell lines as shown by decreased mitochondrial membrane potential and phosphatidylserine externalisation (Figs. 1A and S1B). Potentiation of ABT-737-induced apoptosis in MPM cells was 2DG concentration dependent as illustrated with the MESO-8T cell line (Fig. S1C). Western blotting of numerous BCL-2 family proteins in 2DG-treated MESO-3T,-7T and −8T cells showed that only MCL-1 was downregulated (Fig. 1B). MCL-1 was also down-regulated in MESO-3T cells but with only a small increase in sensitivity at lower concentrations of ABT-737 (Fig. 1A & B). To investigate the specific involvement of MCL-1 in resistant MPM cells, we used MCL-1 siRNA to produce an almost complete MCL-1 knockdown in MESO-7T & 8T cells (Fig. 1C, lanes 3 and 6). BIM expression was also markedly reduced after MCL-1 KD whereas BCL-XL expression decreased only slightly (Fig. 1C, lanes 3 and 6). Knockdown of MCL-1 alone did not induce apoptosis in MESO-7T and MESO-8T cells (Fig. 1D and Fig. S1D). Transfection control or scrambled siRNA cells treated with ABT-737 (10 μM, 4h) showed only a small induction of apoptosis (Fig. 1C, lane 4 & 5; Fig. 1D & Fig. S1D). However, MCL-1 KD markedly enhanced ABT-737 induction of apoptosis (∼90% apoptosis, Fig. 1D, lane 6) which also correlated with complete caspase-9 and PARP cleavage (Fig. 1C. lane 6). These results showed that whilst 2DG potentiated ABT-737-induced apoptosis *via* partial downregulation of MCL-1, complete MCL-1 KD was more potent with ABT-737, inducing nearly complete cell death.

**Figure 1.**
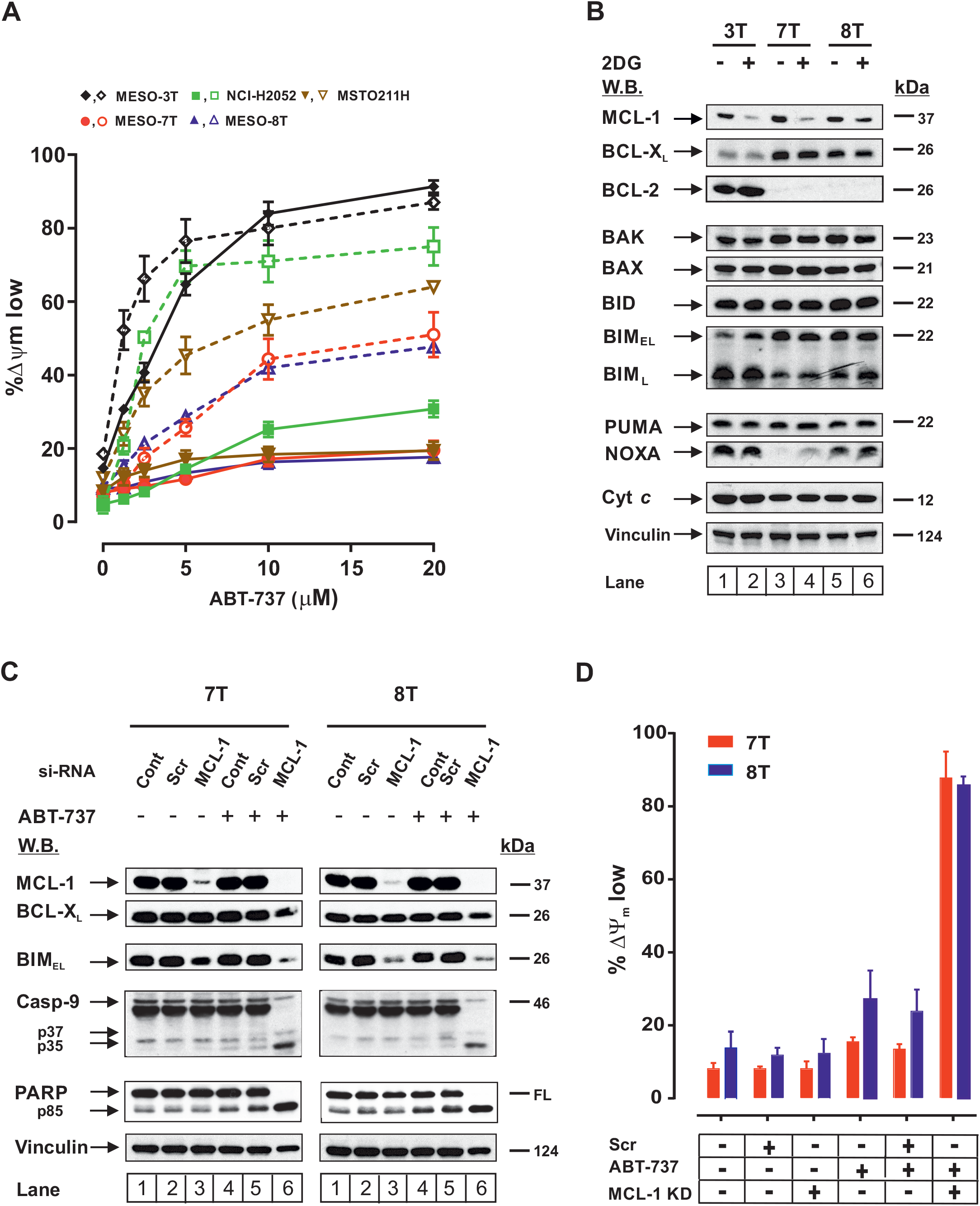
2-deoxyglucose (2DG) potentiates ABT-737-induced apoptosis in MPM cell lines via downregulation of MCL-1. **A)** Primary MESO-3T, −7T and −8T MPM cell lines derived from patients (23) and commercially available cell lines, MSTO-211H and NCI-H2052 were pre-treated with (*open* symbols) or without (*solid* symbols) 2DG (50 mM/16h) followed by ABT-737 (10 μM/6h) at the indicated concentrations. Apoptosis (mean ± SEM, n=4 expts) was assessed by the decrease in mitochondrial membrane potential using TMRE (%ΔΨm low). B) Primary MPM cells treated with 2DG (50 mM/16h) were lysed and cell proteins analysed by SDS-PAGE/Western blotting for the indicated BCL-2 family proteins, Cyt c or Vinculin as a loading control. **C)** Induction of apoptosis in Meso-7T and −8T cells as shown by caspase-9 and PARP cleavage. **D)** MCL-1 siRNA knockdown greatly potentiated ABT-737 (10 μM/6h) induction of apoptosis as assessed by TMRE.

### 2DG-potentiated ABT-737-induced apoptosis is BAX/BAK-dependent but BIM/PUMA independent in primary MPM cells

BH3 mimetics induce apoptosis through BAX/BAK-dependent MOMP and cytochrome *c* release (*30*), so we investigated whether 2DG/ABT-737-induced apoptosis in primary MPM cells was also BAX and/or BAK-dependent. MESO-8T cells with knockdown of either BAX or BAK with siRNA were treated with 2DG or 2DG/ABT-737 (Figs. 2A and 2C & Fig. S2A). Single BAX or BAK KD had no effect on 2DG/ABT-737-induced apoptosis (Fig. 2A, compare lanes 4, 8 & 9). However, BAX/BAK double knockdown (DKD) significantly inhibited 2DG/ABT-737-induced apoptosis (Fig. 2A, compare lanes 4 and 10; Fig. S2A, compare lanes 4 and 10). Similarly, MCL-1 KD/ABT-737-induced apoptosis was not affected by single knockdown of BAX or BAK (Fig. 2B, compare lanes 4, 8 and 9 and Fig. S2B,compare lanes 4, 8 and 9 and Fig. 2C) but was strongly inhibited after BAX/BAK DKD (Figs. 2B and S2B, compare lanes 4 and 10). Thus, 2DG or MCL-1 siRNA knockdown potentiated ABT-737-induced apoptosis was largely dependent on BAX/BAK, but critically both pore-forming BCL-2 family members had to be downregulated for inhibition of cell death.

**Figure 2.**
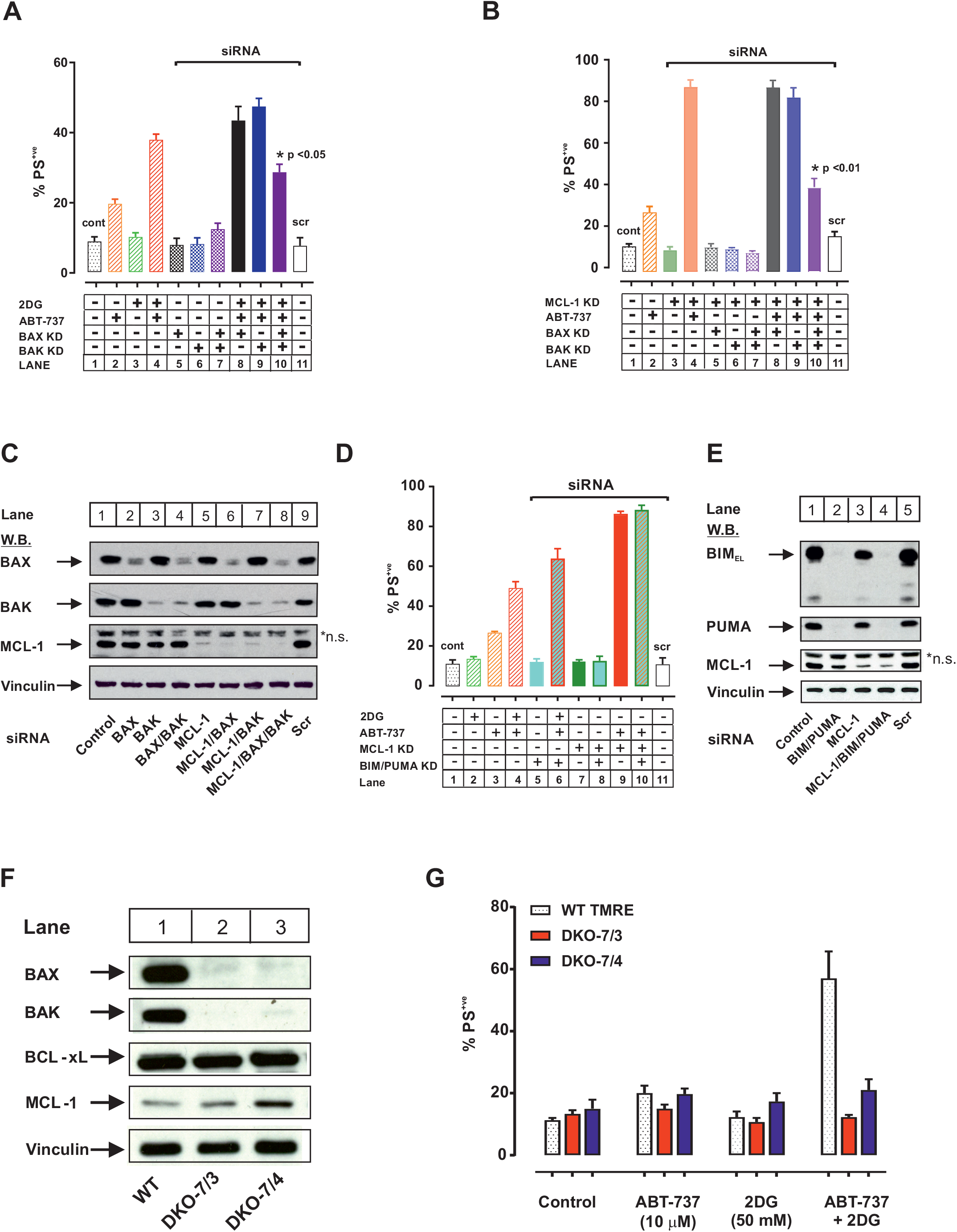
2DG potentiation of ABT-737-induced apoptosis in MPM cells is BAX/BAK dependent, but BIM and PUMA independent. **A**) MESO-8T cells were transiently transfected with BAX and/or BAK siRNAs, followed by 2DG (50 mM/16h) with or without ABT-737 (10 μM/6h) and apoptosis analysed by Annexin-V/PI. **B**) BAX/BAK knockdown cells were also transfected with MCL-1 siRNA,followed by ABT-737 (10 μM/6h) and apoptosis measured by Annexin-V/PI. **C)** Soluble whole cell lysates from Panel B) were analysed for knockdown efficiency of MCL-1, BAX or BAK by immunoblotting. D) MESO-8T cells transiently transfected with BIM and PUMA siRNAs, +/-MCL-1 siRNA or with 2DG prior to ABT-737 (10 μM/6h) and apoptosis assessed by Annexin-V/PI. E) siRNA knockdown of MCL-1 and BIM/PUMA was validated with immunoblotting. F) BAX and BAK were knocked out in MESO-8T cells (DKO-7/3 and DKO-7/4) using CRISPR-Cas9 methodology. G) Potentiation of ABT-737-induced apoptosis in DKO-7/3 and −7/4 cells by 2DG was assessed by Annexin-V/PI assay. Apoptosis analyses (A and B) were from 3 independent experiments (mean ± SEM (n=3), *: p < 0.05, **: p < 0.01, students T-test).

BH3 mimetics can also displace the BH3 activators BIM and/or PUMA from BCL-2/BCL-XL which then directly activate BAX/BAK (*31*). Both BIM and PUMA are highly expressed in the MPM cell lines and using MESO-8T (Fig. 1B) we tested the possible involvement of BIM and PUMA in BAX/BAK activation following 2DG or MCL-1 KD/ABT-737-induced apoptosis (Fig. 2D and E). Double knockdown (DKD) of BIM/PUMA (Fig. 2E) had no impact on either 2DG or MCL-1 KD/ABT-737-induced apoptosis (Fig. 2D, Fig. S2C), which indicated that in MPM cells, BIM and PUMA are not required for 2DG- or MCL-1 KD/ABT-737-induced apoptotic cell death. In our model system, BAX and BAK were significantly, but not totally downregulated by their cognate siRNAs. This may be due to sub 100% transfection efficiency or other factors. Hence, we used CRISPR-Cas 9 to genetically delete BAX and BAK (double knock out, DKO) cells to ensure complete ablation of BAX and BAK in MESO-8T cells (DKO-7/3 & DKO-7/4 clones, Fig. 2F). In these two DKO clones, BAX and BAK could not be detected by Western blotting (Fig. 2F). Importantly, 2DG/ABT-737-induced apoptosis was completely inhibited (Fig. 2G, Fig. S2D). Thus, in ABT-737 resistant mesothelioma cells, inhibition or downregulation of both BCL-2/BCL-XL and MCL-1 allows BAX or BAK to auto activate/oligomerize, resulting in MOMP, cytochrome *c* release, activation of apoptosome and caspases.

### 2DG activates multiple cell signalling pathways which can either antagonise or potentiate ABT-737 cytotoxicity

MCL-1 is a labile protein and has a rapid turnover rate (∼1.5 h) via proteasomal degradation (*32*). Cancer cells have evolved a variety of mechanisms, either by increased protein expression or decreased protein degradation to maintain high expression levels of MCL-1 (*33, 34*). Whilst 2DG is a potent inhibitor of glycolysis it can also potentially effect signalling pathways that may regulate MCL-1 expression (Fig. 3A). To identify these changes we characterised the time-dependent effects of 2DG on these key signalling pathways in MESO-7T/-8T cells (Fig. 3B). 2DG has a biphasic effect inducing a rapid decrease of MCL-1 levels within 2 to 4 h (Fig. 3Bi, lanes 2-3 & 8-9), which remains depressed up to 8 h (Fig. 3 Bi, lanes 4 & 10, Fig. S3A) then partially recovers from 8 to 24 h (lanes 5-6 & 11-12). Thus, 2DG induces an initial rapid decrease in MCL-1, followed by an extended incomplete recovery phase. To investigate this further we characterised the time-dependent activation of key cell survival pathways (Fig. 3A) using phospho-specific antibodies which correlate with their activity status with MCL-1 expression (Fig. 3Bi, Bii and & C). As the RAF/MEK/ERK/STAT3 pathway predominantly regulates transcriptional control of MCL-1 (Fig. 3A) (*33*), STAT3 activity was assayed by immunoblotting for (Y705) p-STAT3. Untreated MESO-7T/8T cells expressed high constitutive levels of p-STAT3 (Fig. 3Bi, lanes 1 & 7), but within 2 h of 2DG treatment, p-STAT3 levels were markedly reduced and from 4-24 h were barely detectable (Fig. 3Bi, lanes 2-6 & 8-12). Total levels of STAT3 were unaffected, whilst its dephosphorylation/inhibition correlated with a ∼ 50% loss of MCL-1 mRNA (Fig. S3A) and 40 % loss of protein (Fig.S3A). STAT-3 Y705 is phosphorylated by ERK 1/2 (Fig. 3A) and in MESO-7T/-8T cells high levels of constitutive p-ERK 1/2 were detected (Fig. 3Bi, lanes 1 & 7) which after 2DG treatment was also rapidly inactivated (2 - 4h), remaining at barely detectable levels at 24 h (Fig. 3Bi). The upstream MEK kinases were also inactivated, although marked inhibition was not apparent until 16 h post treatment (Fig. 3Bi, lanes 5-6 and 11-12). These results are consistent with the possibility that loss of ERK activity following 2DG treatment may not be dependent on the canonical RAS/RAF/MEK pathway and could proceed through another pathway, such as dual-specific phosphatases (DUSP) which are involved in dephosphorylation and subsequent inactivation of MAP kinases (*35*). Strikingly, 2DG also caused rapid inactivation of p38 MAPK, which was sustained up to 24 h post-treatment (Fig. 3Bi).

**Figure 3.**
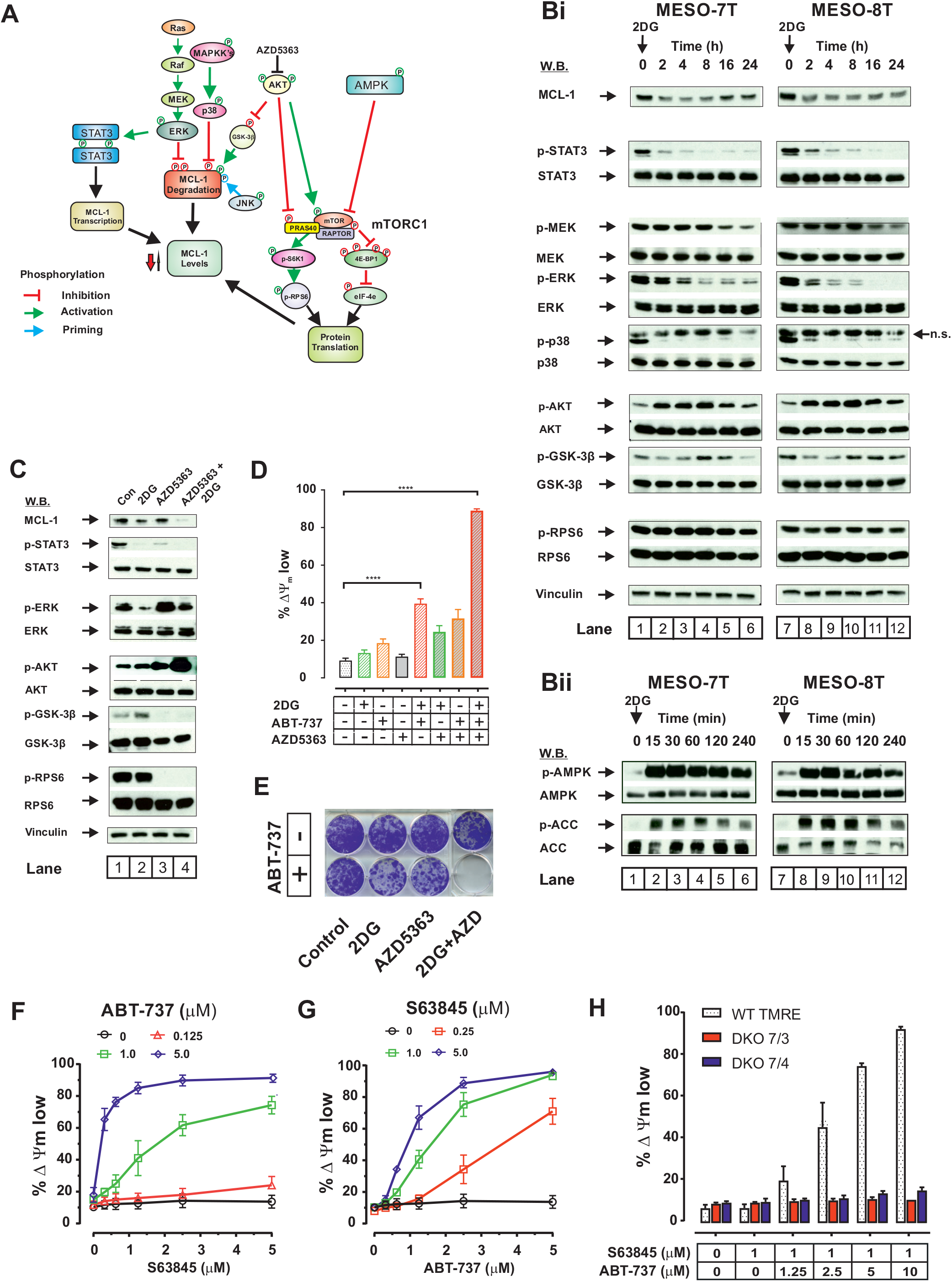
Multiple signalling pathways control 2DG regulation of MCL-1 pro-survival role in mesothelioma cells. A) Schematic showing key signalling pathways potentially involved in regulating MCL-1 protein levels which control its anti-apoptotic function. Bi & Bii) MESO-7T and −8T cells were treated with 2DG (50 mM) for the indicated times and soluble whole cell lysates analysed by immunoblotting. Total and phosphorylated forms of proteins which are signature indicators of the various signalling pathways shown in Figure 3A were analysed. C) MESO-8T cells were treated with or without 2DG, AZD5363 or a combination of 2DG and AZD5363. Immunoblotting profiles for the indicated proteins which map the downstream targets of AKT are shown in lanes 1-4. D) The effects of AZD5363 on 2DG- and ABT-737-induced apoptotic cell death are shown as mean ± SEM (n=4) and statistical analysis performed by one-way ANOVA (**** <0.0001). E) Clonogenic survival was measured with the same combinations of ABT-737, 2DG and AZD5363 as shown in Figure 3D. F) Potentiation of apoptosis (TMRE) in MESO-8T cells with fixed concentrations of ABT-737 (0, 0.125, 1 or 5 μM/ 6 h) and a 0-5 μM concentration range of S63845, a MCL-1 specific inhibitor. G) As in panel F) but with S63845 fixed (0, 0.25, 1 or 5 μM) and ABT-737 varied (0-5 μM). H) Potentiation of apoptosis (TMRE) with 1 μM S6384 (fixed) plus varied ABT-737 (0-10 μM) concentrations in MESO-8T cells is abolished in BAX/BAK DKO cells.

2DG is known to activate AKT in some tumour cells ((*36*) and in our study, we found that AKT activity was increased after 2DG treatment (Fig. 3Bi). Thus, downstream p-GSK3-β was initially dephosphorylated and activated from 0-4 h, followed by a delayed rephosphorylation and inactivation phase, correlating with stabilisation of MCL-1 (Fig. 3Bi, lanes 4 & 5, and 10 & 12). The T163 site of MCL-1 is phosphorylated by p-ERK/p-JNK and believed to be a priming site, leading to phosphorylation by GSK3-β at S155 and S159, targeting MCL-1 for proteasome degradation (*37*). Thus, S159 dephosphorylated GSK3-β is the active kinase and phosphorylation by p-AKT inhibits its activity and limits proteasomal degradation of MCL-1. In both MESO-7T and −8T cell lines, 2DG increased p-AKT levels within 2 h of treatment which peaked at 8 h (Fig. 3Bi, lanes 8 & 10) before decreasing to lower levels by 24 h. Thus, in MESO-7T/8T cells 2DG activates the classic AKT survival-signalling pathway (*38*) in a time-dependent manner which stabilises MCL-1 protein expression (Fig. 3C). In addition, activated AKT can *via* mTOR activation and PRAS40 inhibition activate RPS6 kinase and p-RPS6 to promote enhanced global protein synthesis. Notably, in MESO-7T and −8T cells, p-RPS6 levels were largely maintained throughout 2DG treatment (Fig. 3Bi), whereas ^35^S methionine/cysteine incorporation was reduced by 25 – 40% within 30 min of 2DG treatment (Fig. S3B), indicating that despite active AKT signalling 2DG induced a partial/selective reduction in protein translation.

### Inhibition of AKT markedly enhances 2DG potentiation of ABT-737 cytotoxicity

The AMPK pathway is also targeted by 2DG which produced a rapid decrease in ATP levels (Fig. 4C) and a rapid sustained activation of AMPK (T172) within 15 min correlating with the kinetics of phosphorylation of acetyl-CoA carboxylase (ACC), an AMPK substrate (Fig. 3Bii). Activated AMPK inhibited mTOR and its downstream targets 4E-BP1 and eIF-4e (*39*), which partly inhibited protein translation as shown with the ^35^S incorporation data (Fig. S3B). Thus, the rapid effect of 2DG on AMPK activation which may inhibit protein translation is potentially antagonised by the delayed activation of the AKT pathway. Consequently, we hypothesized that an AKT inhibitor could potentiate the effects of 2DG on MCL-1 downregulation and consequent ABT-737-induced cell death. AZD5363 is an AKT inhibitor currently in Phase II clinical trials for treating breast and non-small cell cancers (*40*) and is therefore a potentially clinically relevant drug to pair with 2DG. Thus, 2DG/16 h treatment significantly decreased p-ERK/p-STAT3 and MCL-1 levels with little effect on p-RPS6 (Fig. 3C, lane 2). In contrast, AZD5363 on its own increased p-ERK and ablated STAT3 and RPS6 phosphorylation, without markedly affecting MCL-1. AZD5363 also completely abolished GSK3-β phosphorylation, coupled to increased p-AKT (S473) levels. AKT phosphorylation following AZD5363 treatment can be explained by the fact that p-S6K1 is a downstream target of p-AKT/mTORC1 and a negative regulator of mTORC2 and IRS-1 which regulate AKT phosphorylation and activation (Fig. 3A). Inhibiting p-AKT activity leads to dephosphorylation of p-S6K1 (as shown by the loss of p-RPS6, Fig. 3C), releasing IRS-1 to recruit PI3K to the plasma membrane thus activating AKT *via* PDK1 phosphorylation of T308 and mTORC2 phosphorylation of AKT S473 (*41*). In contrast, the combination of 2DG and AZD5363 resulted in hyper-phosphorylation of AKT and total ablation of MCL-1 and p-RPS6. AZD5363 alone did not induce cell death in MESO-8T cells (Fig. 3D), but combined with ABT-737 resulted in ∼ 35% cell killing. 2DG alone as shown above (Fig. 1) did not reduce cell viability but did enhance the effect of both ABT-737 (40%) and AZD5363 (30%)-induced cell killing. Significantly the triple AZD5363/ABT-737/2DG combination was lethal, producing ∼90% cell death (Fig. 3D). Thus, AZD5363/2DG enhances loss of MCL-1 and also abrogates protein translation, resulting in a complete loss of MCL-1. Under these conditions ABT-737 targets BCL-2 and produces ∼90 % cell killing and also completely ablates clonogenic cell growth (Fig. 3D & 3E).

**Figure 4.**
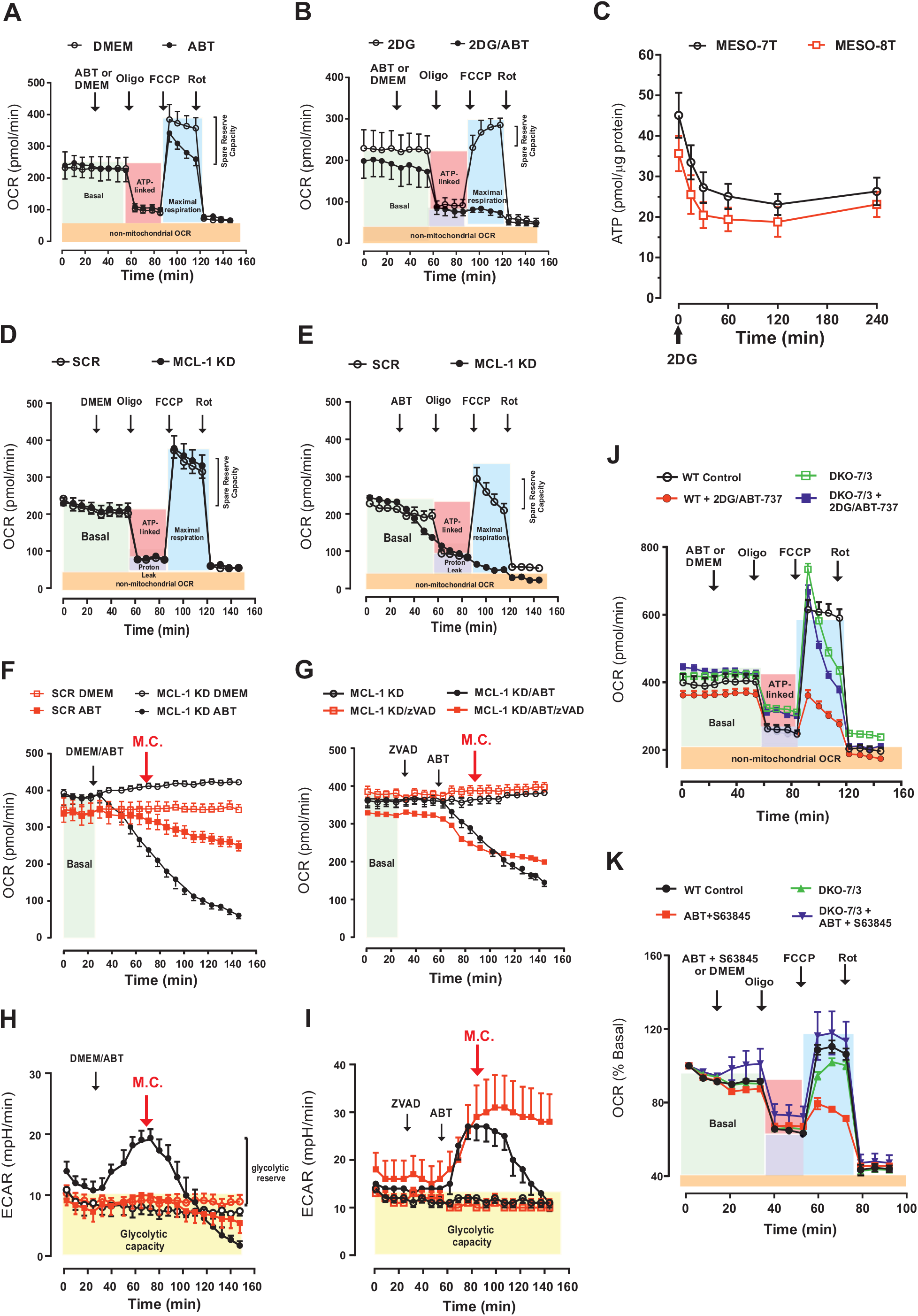
ABT-737 inhibits OXPHOS and induces Caspase-dependent inhibition of glycolysis in MCL-1 knockdown cells. OXPHOS (OCR) and glycolysis (ECAR) of MPM cells were analysed in Seahorse XF24/96 extracellular bioanalysers. A) The mitochondrial OXPHOS stress test was performed on Meso-8T cells with DMEM or ABT (ABT-737, 10 μM), followed by Oligo (Oligomycin, 400 nM), FCCP (400 nM) and Rot (1 μM Rotenone). B) As in panel A, except that cells were pre-treated with 2DG (50 mM/4h). C) 2DG was added to cells and a time-dependent decrease in ATP levels measured. D and E) Similar to panels A and B, except that the cells were transfected with MCL-1 or a scrambled (SCR) control siRNA. F) Effect of ABT-737 on OCR in MCL-1 KD cells and corresponding stimulation of ECAR (panel H). M.C.indicates the onset of a metabolic catastrophe with complete loss of glycolytic activity (M.C), accompanied by decreasing OCR. In contrast, inhibition of OCR by oligomycin and rotenone also induced a rapid increase in ECAR but unlike MCL-1/ABT-737 this was sustained (Figure S5A). G and I) As in Panels F and H respectively, except cells were pretreated with zVAD.fmk, a pan-caspase inhibitor, which did not prevent the inhibition of OCR, but blocked the M.C., and resulted in a sustained elevation of ECAR activity. J) The experiments with 2DG and ABT-737 in panel (B) were repeated in BAX/BAK DKO-7/3 cells showing that BAX/BAK deletion blocked the effects of 2DG and ABT-737. K) Similar experiments performed with BAX/BAK double knockout cells treated with ABT-737 in the presence of S63845.

The above results show that the combination of 2DG and simultaneously targeting the AKT pathway can overcome the resistance and cell survival mechanisms operating in mesothelioma cells. This strategy is certainly effective but is targeting multiple cell signalling pathways, so we aimed to simplify the combinatorial approach by subsequently specifically targeting MCL-1 which underpins the resistance of MPM cells to ABT-737-induced apoptosis. To this end we used the MCL-1 inhibitor, S63845, which greatly increased both the sensitivity and extent of apoptosis (∼90%) induced by ABT-737 in MESO-8T cells (Fig. 3F & G). This potentiation of cell killing was also BAX/BAK-dependent, as BAK/BAK DKO cells (DKO-7/3 & 7/4) failed to undergo ABT-737/S63845-induced apoptosis (Fig. 3H and Fig. S3C). These results provide a mechanistic rationale for successfully targeting and overcoming MPM tumour cell resistance.

### Targeting BCL-XL and MCL-1 inhibits both OXPHOS and glycolysis and induces a metabolic catastrophe

2DG is actively transported into tumour cells *via* the glucose transporter GLUT1 which mediates the increased demand for glucose (*6*). Hexokinase (HK) converts 2DG into 2DG-6-phosphate, which is not metabolised by the glycolytic pathway, and competitively inhibits HK and non-competitively inhibits phosphoglucoisomerase (PGI), which are the critical first steps in glycolysis. As 2DG is a potent glycolytic inhibitor it potentially switches the cell to mitochondrial oxidative phosphorylation (OXPHOS) and we therefore investigated the effects of 2DG on the bioenergetic capacity of MPM cells and the possible impact on the mitochondrial apoptotic pathway. OXPHOS and glycolysis were assessed simultaneously by measuring oxygen consumption rate (OCR) and lactate release (ECAR) respectively with Seahorse XF24/96 analysers (*18*). Typically, MESO-7T/-8T cells had basal OCRs of ∼175-230 pmol/min/ 5 × 10^4^ cells (Fig. 4A & Fig. S4A) and ECARs of ∼ 5-6 mpH/min (Fig. S4C). Pre-treatment with 50 mM 2DG/4 h resulted in small increases in OCR (Fig. 4B & Fig. S4B) and a marked reduction in ECAR (<2 mpH/min, Fig. S4D), consistent with a bioenergetic switch from glycolysis to OXPHOS. ATP levels rapidly (0-30 min) decreased by ∼50% and then plateaued from 30-240 min (Fig. 4C). MESO-7T/-8T cells responded to a classical mitochondrial ‘stress test’ using oligomycin (ATP synthase inhibitor), FCCP (uncoupler) and rotenone (ETC inhibitor). Oligomycin inhibited basal OCR (∼ 240 to ∼ 90 pmol/min, Fig. 4A), and after adjusting for non-mitochondrial OCR (∼40 pmol/min) indicates that ∼75% of basal OCR is linked to ATP production. Inhibition of ATP-linked OCR was accompanied by a 2-fold compensatory increase in ECAR (∼11 mpH/min, (Fig. S4D). The increase in ECAR (‘glycolytic reserve’) was abolished by 2DG (Fig. S4F) and the FCCP stimulated (maximal) respiration was reduced in 2DG treated cells along with a decrease in the ‘spare respiratory capacity’ (compare Fig. 4B and S4C). Without 2DG, ABT-737 did not affect basal OCR and ECAR levels (Fig. 4A & Fig. S4E) and the response to oligomycin, but did slightly inhibit FCCP-stimulated respiration (Fig. 4A). In the presence of 2DG (50 mM/4 h), basal OCR and ATP-linked respiration were not affected by ABT-737 (Fig. 4B), but FCCP-stimulated respiration was now almost completely inhibited (Fig. 4B). Thus, 2DG inhibition of glycolysis switches MPM cells to OXPHOS, decreasing ‘spare respiratory capacity’ and this metabolic stress sensitizes the cells to ABT-737.

As 2DG/ABT-737-potentiated cell death is dependent on MCL-1 down-regulation, and is accompanied by inhibition of OXPHOS, we explored the possible role of MCL-1 in cellular bioenergetics using two approaches, MCL-1 depletion and S63845-mediated inhibition. Significantly, in MPM cells, MCL-1 KD alone did not affect OXPHOS (Fig. 4D), but did potentiate the effects of ABT-737, which now completely inhibited both basal and FCCP-stimulated (maximal) OCR (Fig. 4E). ABT-737 in control cells induced a time-dependent (30-150 min) slow but small decrease in basal OCR (Fig. 4F) without affecting ECAR (Fig. 4H). However, in MCL-1 KD cells, ABT-737 induced an extensive and rapid inhibition of basal OCR (Fig. 4F), which correlated with an immediate (∼ two fold) increase in ECAR, which peaked at 70 min before declining to zero at 150 min (Fig. 4 H). This bell shaped curve indicated that in MCL-1 KD cells, ABT-737 is a potent OXPHOS inhibitor and induces a transitory compensatory increase in ECAR which is not sustained, resulting in a subsequent “Metabolic Catastrophe” (M.C.). The canonical OXPHOS inhibitors oligomycin and rotenone also rapidly inhibited basal OCR and induced a compensatory increase in ECAR, which in contrast to ABT-737, was sustained without a M.C (Fig. S5A & B). However, ABT-737, unlike rotenone and oligomycin, in MCL-1 KD cells induces apoptosis *via* caspase activation (Fig. 1C). Notably, pre-treatment with the pan-caspase inhibitor Z-VAD.fmk prior to ABT-737 did not prevent the decrease in OCR (Fig. 4G) showing that the OXPHOS blockade was caspase-independent. While Z-VAD.fmk did not block the ABT-737-induced initial increase in ECAR, Z-VAD sustained this increase in ECAR, importantly without a M.C. (Fig. 4I), phenocopying the effects observed with rotenone or oligomycin (Fig. S5A and B).

BAX/BAK deletion using CRISPR-Cas9 gene editing, which blocked ABT-737/2DG-induced apoptosis (Fig. 2F, 2G and S2D) in DKO-7/3 cells, had little effect on cellular bioenergetics *per se* (Fig. 4J). 2DG did not affect the ‘stress test’ response of the DKO-7/3 cells, although interestingly the FCCP-stimulated OCR was not sustained (Fig. 4J). In the presence of 2DG, ABT-737 inhibited FCCP-stimulated OCR in WT cells (similar to Fig. 4B) but not in the DKO-7/3 cells showing that ABT-737 inhibition of maximal respiration was BAX/BAK–dependent. Similarly, using S63845 to directly target MCL-1 in combination with ABT-737 inhibited FCCP-stimulated respiration in wild type cells, but was ineffective in DKO-7/3 cells (Fig. 4K). Thus, if MCL-1 is deleted or inhibited then ABT-737 induces MOMP resulting in cytochrome *c* release and inhibition of the ETC. Strikingly, we observed two key glycolytic enzymes, phosphofructokinase and pyruvate kinase were also cleaved after S63845/ABT-737 treatment and this cleavage was abolished with Z-VAD.fmk, under experimental conditions where S63845/ABT-737-induced pro-caspase-9 cleavage/activation was also blocked by Z-VAD.fmk (Fig. S5C). Of note, Hexokinase I (HK I), another critical glycolytic enzyme was not cleaved upon S63845/ABT-737 treatment, indicating that caspase-mediated destruction of glycolytic enzymes is both selective and specific (Fig. S5C). The combination of S63845 and ABT-737 induced a time-dependent release of mitochondrial cytochrome *c* and Smac into the cytosol (Fig. S5D) which is consistent with MOMP activation, apoptosome formation and apoptosis induction. Thus, the glycolytic collapse induced by ABT-737 and MCL-1 KD or MCL-inhibitor is a caspase-dependent event which results in cleavage of glycolytic enzymes which disables the glycolytic pathway (M.C.) and enhances the cell killing effect of ABT-737.

### Targeting MCL-1 sensitises mesothelioma patient-derived 3D tumour explants to ABT-737-induced apoptosis

Currently, the therapeutic efficacy of potential antitumor agents is largely evaluated in 2D cell culture systems. We therefore wished to determine whether the combinatorial treatments described above could be equally effective in a more clinically-relevant patient-derived 3D explant model. The mechanisms involving the spatiotemporal evolution of tumour development are reproduced in our 3D model, which more closely recapitulates the entire tumour microenvironment. Explants from freshly dissected tumours were layered on top of tissue culture inserts (Fig. S6). Explants were treated with various drugs prior to fixation and then processed for quantitative immunohistochemical analysis (Fig. S6 and Materials and Methods). Down regulation of MCL-1 in cytokeratin-positive (MPM tumour) cells (*h*-score) was observed in either 2DG-treated samples or in explants treated with 2DG and AZD5363 (Fig. 5A & B, lanes 2, 5, 7 and 8). Apoptosis in mesothelioma cells was assessed by overlapping cytokeratin-positive cells with cleaved PARP-positive cells and normalized against individual patient’s control (Fig. 5C). 2DG alone induced a small, but significant increase in apoptosis (Fig. 5C, lane 2) and significantly enhanced ABT-737-induced apoptosis (Fig. 3C, lane 5 p<0.05). Importantly, tumour areas which exhibited apoptosis were correlated with those areas in which MCL-1 levels were decreased (Fig. 5A, B and C). These results showed that 2DG decreased MCL-1 levels and potentiated ABT-737-induced apoptosis in not only primary MPM cell lines but also in the 3-D model system. AZD5363 on its own also induced a small increase in apoptosis which was slightly enhanced by 2DG treatment (Fig. 5C, lanes 4 and 7). Importantly, as in the 2D primary cell experiments, the triple combination of ABT-737/2DG/AZD5363 was the most potent treatment leading to an 8-fold increase in apoptosis in the cytokeratin-positive cells (Figs. 5A and C, lane 8). Finally, the induction of apoptosis by ABT-737 in cytokeratin-positive cells in the explant model was potentiated ∼ 8-fold by inhibiting MCL-1 with S63845 (Fig. 5D & 5E) with significantly less effect on the surrounding tumour stroma cells (data not shown).

**Figure 5.**
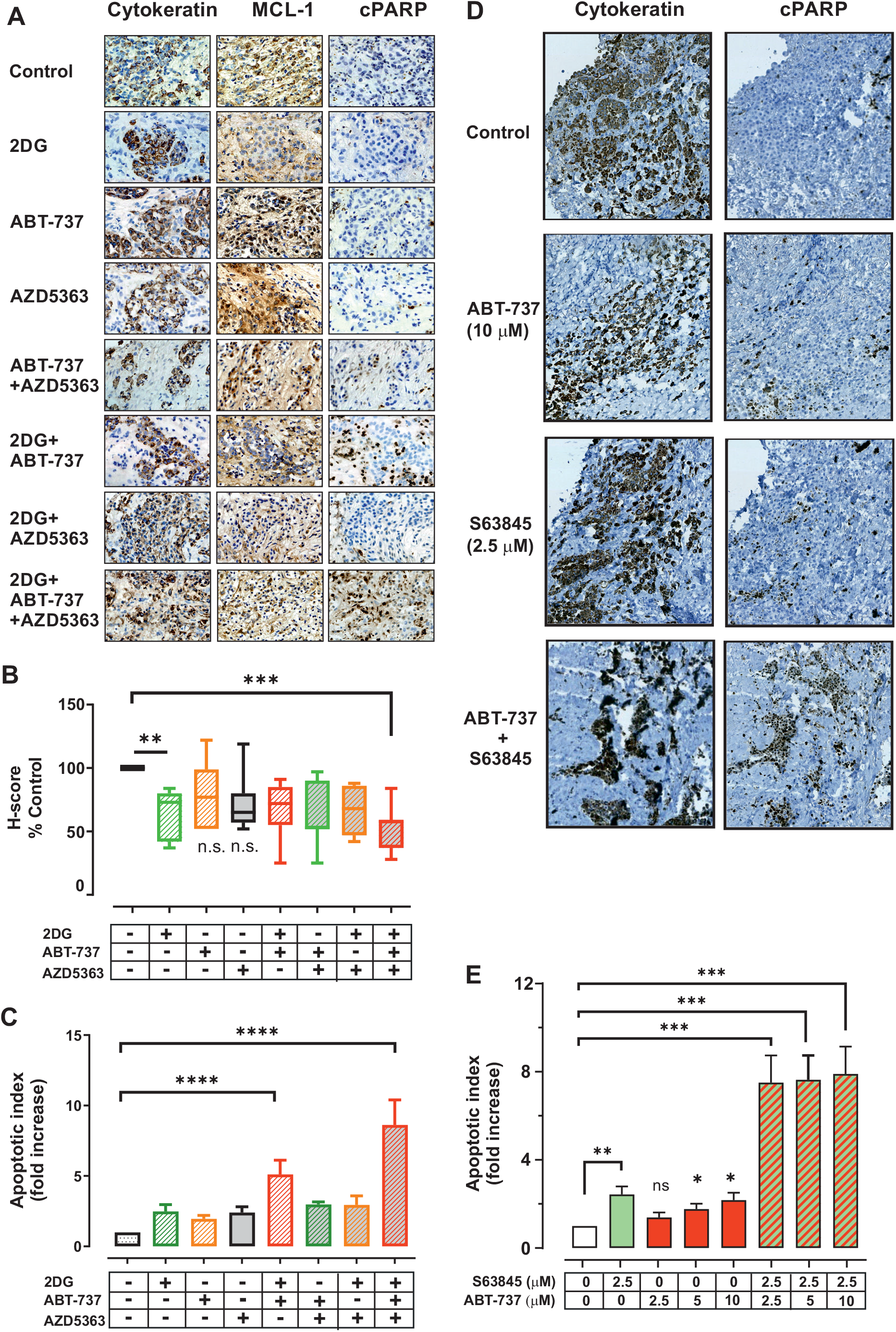
Targeting MCL-1 protein expression sensitizes MPM cells in mesothelioma explants to ABT-737-induced apoptosis. Tissue (2 mm^3^) cubes were cut from mesothelioma tumour samples obtained from surgical operations with patient consent and positioned on organ culture inserts (Figure S6A). Tissues cubes were incubated for 24 h with either culture medium only or media containing either 2DG (50 mM), ABT-737 (10 μM), AZD5363 (20 μM) or combinations of ABT-737 + AZD5363, 2DG + ABT-737, 2DG + AZD5363 and 2DG + ABT-737 + AZD5363. Treated tissues were fixed and processed using conventional histological procedures, followed by immunohistochemistry as described in the methods using primary antibodies to cytokeratin (MPM tumour marker), MCL-1 and cleaved PARP (cPARP, apoptosis marker). B & C) Adjacent slides were analysed with Visiopharm software to quantify (B) MCL-1 levels and (C) the percentage of cleaved PARP (cPARP) in MPM (cytokeratin-positive) cells. Data were normalised within individual patient samples to use control cells as 100 % for MCL-1 expression and 1 for the apoptotic index. Data from 7 patients shown as mean ± SEM, **: p < 0.001, ***: p < 0.0001 and statistical analysis by one-way Anova. D) and E, Effect of ABT-737 and S63845 on cPARP expression in cytokeratin-positive cells. E) Potentiation of ABT-737-induced apoptosis by S63845 (2.5 μM) in explant tissue with increasing concentrations (2.5, 5 and 10 μM) of ABT-737. Apoptotic index calculated from 10 patient samples and statistical analysis by one-way Anova.as shown in panel C and expressed as mean ± SEM; ****: p < 0.0001.

## Discussion

Bioenergetic changes (Warburg effect) and dysregulation of the BCL-2 family of proteins are regarded as hallmarks and strong drivers of cancer growth and resistance to chemotherapy (*42*). In this study, we have co-targeted these two hallmark drivers to induce cell killing in clinically relevant patient-derived MPM 2D and 3D model systems. We show that ABT-737, which inhibits both BCL-2 and BCL-XL, is not a potent inducer of apoptosis in the MPM cell lines or explants (Fig. 1 & 5). Similarly, 2DG which ablates the glycolytic pathway, does not significantly induce cell killing in either MPM cell lines (Figs. 1 and 2) or in 3D explants (Fig. 5). In marked contrast 2DG significantly enhances ABT-737-induced cell killing in both patient-derived 2D and 3D MPM model systems (Figs. 1A, 2A and 5). 2DG has been shown to inhibit aerobic glycolysis in many tumor cells (*6, 8, 19, 43*) and to sensitize these cells to apoptotic stimuli, especially those agents which target the anti-apoptotic proteins BCL-2/BCL-XL/MCL-1. The main effect of 2DG in many cancer cells is to downregulate MCL-1 protein expression and we show for the first time this also occurs in primary patient-derived 2D MPM cell lines and of particular note, 3D explants (Figs. 1B and 5). In all the patient derived cells and immortalized cell lines we have profiled (Fig. S1), MCL-1 is consistently expressed along with BCL-XL. However, BCL-2 is absent from most of these cell lines, except for primary cells MESO-3T, −12T and −50T and the MSTO-211H cell line. Interestingly, MESO-3T cells in comparison to the MESO-7T/-8T are essentially partially sensitive to ABT-737 on its own (50% apoptosis with ∼ 1 μM ABT-737), presumably due to relatively low expression of MCL-1.

We also show that the downregulation of MCL-1 by 2DG is *via* its effects on multiple cell signaling pathways, which combine to block protein transcription/translation (Fig. 3). We have also shown that this enhanced cell killing requires BAX and BAK, but not BIM, PUMA or NOXA and essentially involves BCL-XL and MCL-1 which both bind to BAX and BAK to stop MOMP, cytochrome *c* release and apoptosis. In resistant MPM cell lines and explants, 2DG enhanced ABT-737 killing efficiency but did produce total cell killing (40-50% in MESO-7T & 8T) and this may be due to the fact that the 2DG effect on MCL-1 total protein levels is antagonized by co-activation of the AKT pathway, as the combination of 2DG and the Akt inhibitor AZD5363 resulted in total loss of MCL-1 compared to 2DG only (Fig. 3C). This acts as a survival pathway by inhibiting MCL-1 degradation, partially restoring MCL-1 levels (Fig. 3Bi). In support of this hypothesis, inhibition of AKT activity with AZD5363 potentiated ABT-737 cell killing and in combination with 2DG resulted in complete loss of MCL-1 in MPM cells, ∼90% apoptosis and total blockade of clonogenic cell growth (Fig. 3C, 3D & 3E) and importantly an 8-fold increase in apoptotic cell death cytokeratin-positive (tumour) cells in Explants (Fig. 5). The importance of MCL-1 in protecting MPM cells from ABT-737-induced cell killing was further highlighted by the MCL-1 siRNA experiments showing that MCL-1 KD resulted in enhanced total cell killing with ABT-737 (Fig. 2B, ∼85-95%). Furthermore, the combination of S63854 and ABT-737 was similarly potent in inducing apoptosis in both MPM 2D cell lines and 3D explants (Figs. 3F and 3G, Fig. 5).

In our studies, both the primary MPM cell lines and the *ex vivo* 3D explants were obtained from MPM patients, enhancing the clinical application of our study. Our pre-clinical findings are thus very encouraging as they show that by directly targeting and nullifying the anti-apoptotic MCL-1 protein, it renders intractable MPM cells to be successfully targeted by inhibiting BCL-XL. Our study shows that MPM cannot be successfully treated with either a BCL-2/BCL-XL or MCL-1 inhibitor alone and some rational combinatorial regimes will be needed to target this hitherto untreatable disease. Linking together the bioenergetic and anti-BCL-2 family hallmarks of MPM can be achieved by using 2DG/ABT-737 which certainly enhances the sensitivity of MPM cells to apoptotic death by partially but not totally downregulating MCL-1 protein levels. Using MCL-1 KD or an AKT inhibitor to enhance the effect of 2DG results in total MCL-1 loss and correlates with essentially 100% MPM cell killing. MCL-1 loss renders OXPHOS in MPM cells vulnerable to ABT-737 by loss of mitochondrial cytochrome *c* which inhibits the ETC and respiration. The loss/release of cytochrome *c* has another advantage in that it activates the apoptosome and caspase cascade, resulting in caspase-dependent cleavage of glycolytic enzymes and subsequent inhibition of glycolysis (Fig. S5C). This is in effect a Metabolic Catastrophe as both ATP (energy) producing pathways are disabled, which will enhance cell killing in tumor cells. The triple 2DG/ABT-737/AZD5363 combination is very effective against MPM. The combination of ABT-737 and the MCL-1 inhibitor, S63458,, is also equally effective at killing MPM cells and mirrors other studies with leukemic cells using venetoclax/ABT-199 and S63458 (*44*). There are now a number of MCL-1 inhibitors under development and some have advanced to clinical trials (for review see (*45*)), although one trial with AMG397 an oral MCL-1 inhibitor has been halted due to concerns about cardiotoxicity (). In this respect it may be that using 2DG to enhance ABT-737 cell killing could be more tumor specific as the enhanced uptake of 2DG into the tumor cell will result in a more targeted inhibition of BCL-2/BCL-XL and subsequent induction of apoptosis in MPM cells.

## Supplemental Figure Legends

**Figure S1** - A) BCL-2/BCL-XL/MCL-1 expression levels in diverse patient derived MPM (MESO-*T) primary cells and commercial cell lines. B) Apoptotic cell death in MESO-3T, - 7T, 8T primary cells and NCI-H2052, MSTO211H cell lines treated with ABT-737 alone (*solid* symbols) or with 2DG (*open* symbols) assayed by Annexin V/PI (% PS^+ve^) assay (mean ± SEM, n=4). C) 2DG concentration-dependent potentiation of ABT-737 induction of apoptosis assayed by decreased mitochondrial membrane potential (%ΔΨm low) using TMRE (mean ± SEM, n=4). D) % PS^+ve^ assay (Annexin V/PI) for induction for apoptosis with ABT-737 in MESO-7T & 8T cells after siRNA MCL-1 KD.

**Figure S2** - A) TMRE assay for induction of apoptosis with ABT-737 and 2DG after BAX/BAK siRNA knockdown. B) TMRE assay for induction of apoptosis with ABT-737 after BAX/BAK/MCL-1 siRNA knockdown. C) TMRE assay for induction of apoptosis with ABT-737 after MCL-1/BIM/PUMA KD. D). TMRE assay for induction of apoptosis with ABT-737 and 2DG in MESO-8T WT and BAX/BAK DKO-7/3 and 7/4 clones.

**Figure S3** - A) MCL-1 cellular protein and mRNA levels assayed by immunoblotting and RT-PCR after 2DG (50 mM) treatment of MESO-8T cells. B) Protein (^35^S labelling) synthesis in MESO-7T & −8T cells following 2DG (50 mM) treatment for 0-8h. C) Annexin V/PI assay for induction of apoptosis with S63845 + ABT-737 in MESO-8T WT, BAX/BAK DKO-7/3 and 7/4 clones.

**Figure S4** - A) OXPHOS (OCR) profiles in MESO-7T & 8T cells after mitochondrial OXPHOS stress test using oligomycin (Oligo), FCCP and rotenone (Rot). B) OCR after mitochondrial stress test in MESO-7T & 8T cells after 2DG pre-treatment. C). Effect on glycolysis (ECAR) after mitochondrial stress test using Oligo, FCCP and Rot followed by 2DG (50 mM). D) Effect on glycolysis (ECAR) after 2DG (50 mM) followed by mitochondrial stress test. E) Effect of ABT-737 on glycolysis (ECAR), followed by mitochondrial stress test. F) Effect of ABT-737 on glycolysis (ECAR) in cells pre-treated with 2DG (50mM) followed by mitochondrial stress test.

**Figure S5** - A) Effect of oligomycin or rotenone on OXPHOS (OCR) in MESO-8T cells after si-RNA MCL-1 KD. B) Effect of oligomycin or rotenone on glycolysis (ECAR) in MESO-8T cells after si-RNA MCL-1 KD. C) Caspase-dependent cleavage of pro-caspase-9, pyruvate kinase (PKM2) and phosphofructokinase (PFKP) after induction of apoptosis with S63845/ABT-737 following pretreatment with or without Z-VAD.fmk. D) Immunoblotting showing time-dependent cytochrome *c* and Smac release from mitochondria (P) into the supernatant (S) after S63845 and ABT-737 treatment in a

**Figure S6** - A) Methodology of explant experiment. B) Example of induction of apoptosis as detected by immunohistochemistry using cleaved PARP (cPARP) in cytokeratin positive (tumour) cells after ABT-737+S63845, cPARP/Cytokeratin.

**SUPPLEMENTAL TABLE 1.**

| REAGENT or RESOURCE | SOURCE | IDENTIFIER |
| --- | --- | --- |
| Antibodies |  |  |
| Rabbit polyclonal MCL-1 | Santa Cruz | Sc-819 |
| Mouse monoclonal BCL-2 | Santa Cruz | Sc-509 |
| Rabbit polyclonal BCL-XL | BD | 610212 |
| Rabbit polyclonal BAX | Millipore | ABC11 |
| Rabbit polyclonal BAK | Millipore | 06-536 |
| Rabbit polyclonal BID | Santa Cruz | Sc-11423 |
| Rabbit polyclonal BIM | Millipore | 202000 |
| Rabbit polyclonal PUMA | Cell Signalling Technology | 4976 |
| Mouse monoclonal NOXA | Millipore | OP-180 |
| Mouse monoclonal cytochrome c | Pharmingen | 556433 |
| Rabbit polyclonal caspase-9 | this group | Sun et al., 1999 |
| Mouse monoclonal vinculin | Abcam | Ab18058 |
| Mouse monoclonal PARP | ENZO | BML-SA249 |
| Rabbit polyclonal phospho ERK | Cell signalling Technology | 9101 |
| Rabbit monoclonal ERK | Cell Signalling Technology | 4695 |
| Rabbit polyclonal phospho P38 | Cell Signalling Technology | 9211 |
| Rabbit polyclonal P38 | Cell Signalling Technology | 9212 |
| Rabbit monoclonal phospho STAT3 | Abcam | Ab76315 |
| Rabbit monoclonal STAT3 | Cell Signalling Technology | 4904 |
| Mouse monoclonal phospho AKT | Cell Signalling Technology | 4051 |
| Rabbit monoclonal AKT | Cell Signalling Technology | 4691 |
| Rabbit monoclonal phospho GSK3β | Cell Signalling Technology | 9323 |
| Rabbit monoclonal GSK3β | Cell Signalling Technology | 9315 |
| Rabbit monoclonal phospho AMPK | Cell Signalling Technology | 2535 |
| Rabbit monoclonal AMPK | Cell Signalling Technology | 2603 |
| Rabbit polyclonal phospho S6RP | Cell Signalling Technology | 2215 |
| Rabbit monoclonal S6RP | Cell Signalling Technology | 2217 |
| Rabbit monoclonal phospho MEK | Cell Signalling technology | 9154 |
| Mouse monoclonal MEK | Cell Signalling Technology | 4694 |
| Rabbit monoclonal phospho ACC | Cell Signalling Technology | 11818 |
| Rabbit monoclonal ACC | Cell Signalling Technology | 3676 |
| Mouse monoclonal cytokeratin | DAKO | M0821 |
| Rabbit monoclonal PFKP | Cell Signalling Technology | 5412 |
| Rabbit monoclonal PKM2 | Cell Signalling Technology | 4053 |
| Rabbit monoclonal HKII | Cell Signalling Technology | 2867 |
| Rabbit polyclonal Smac | made in this lab | Sun et al., 2004 |
| Rabbit cleaved PARP | Abam | ab32064 |
| Chemicals |  |  |
| 2-deoxyglucose (2DG) | Sigma | D8375 |
| ABT-737 | Selleckchem | S1002 |
| AZD5363 | Selleckchem | S8019 |
| Oligomycin | Sigma | O4876 |
| FCCP | Sigma | C2920 |
| Rotenone | Sigma | R8875 |
| ZVAD-fmk | MP Biomedicals | FK109 |
| EasyTag35S Protein Labeling Mix | PerkinElmer | NEG772002MC |
| ULTIMA-FLO (scintillation Fluid) | PerkinElmer | 6013589 |
| TMRE | Cambridge Biosciences | BT70016 |
| Annexin V-FITC | This lab | Kohlhaas et al., 2007 |
| RNAiMAX | Invitrogen | 13778 |
| Cell-Titer-Glo Kit | Promega | G7570 |
| Passive Lysis Buffer | Promega | E1941 |
| TRI-Reagent | Sigma | T9424 |
| Superscript III | Life Technologies | 18080093 |

**SUPPLEMENTAL TABLE 2.**
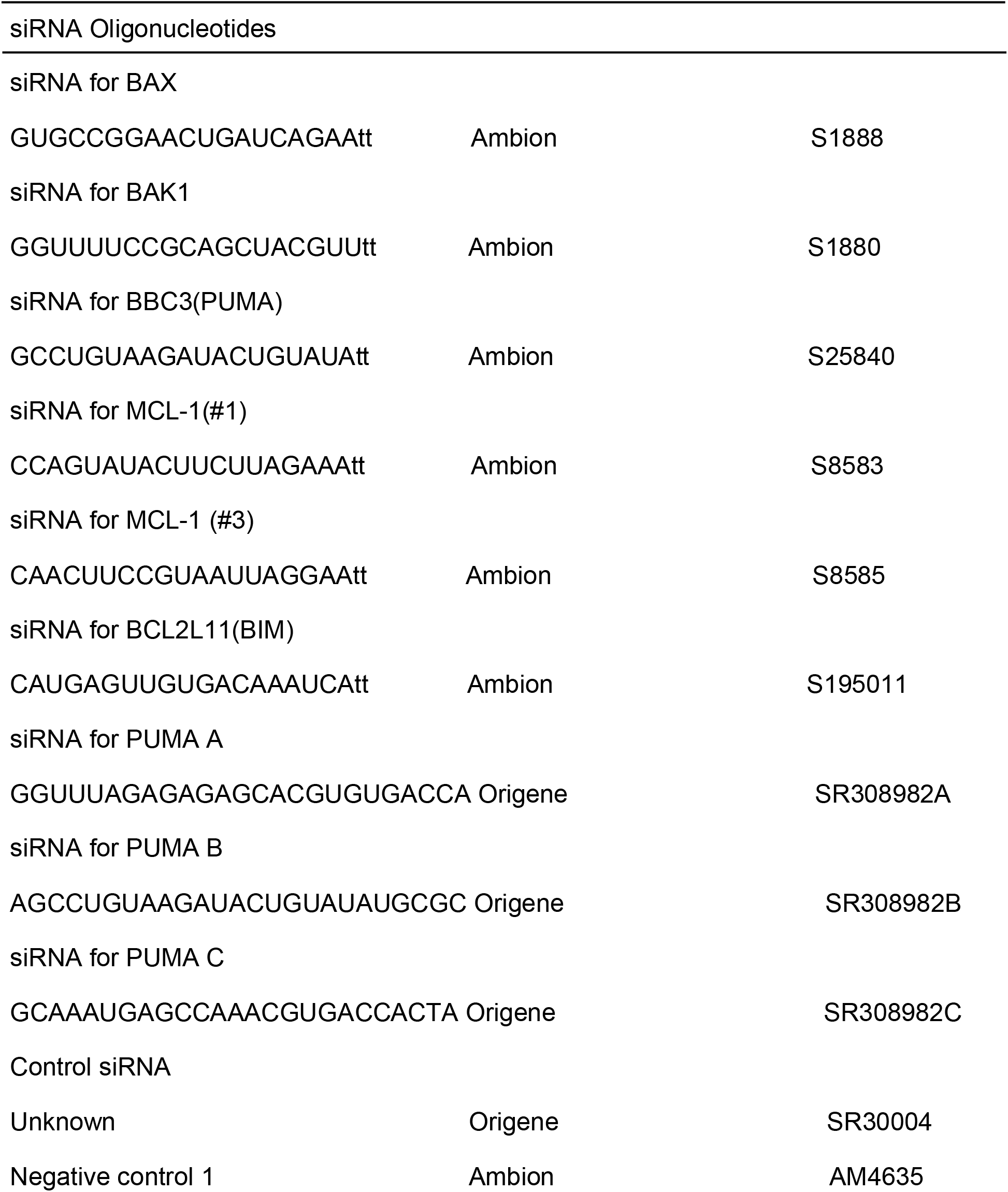

