## Supplementary figures and images for "Metabolic reprogramming provides a novel approach to overcome resistance to BH3-mimetics in Malignant Pleural Mesothelioma"

### Supplemental Figures_S1-S6

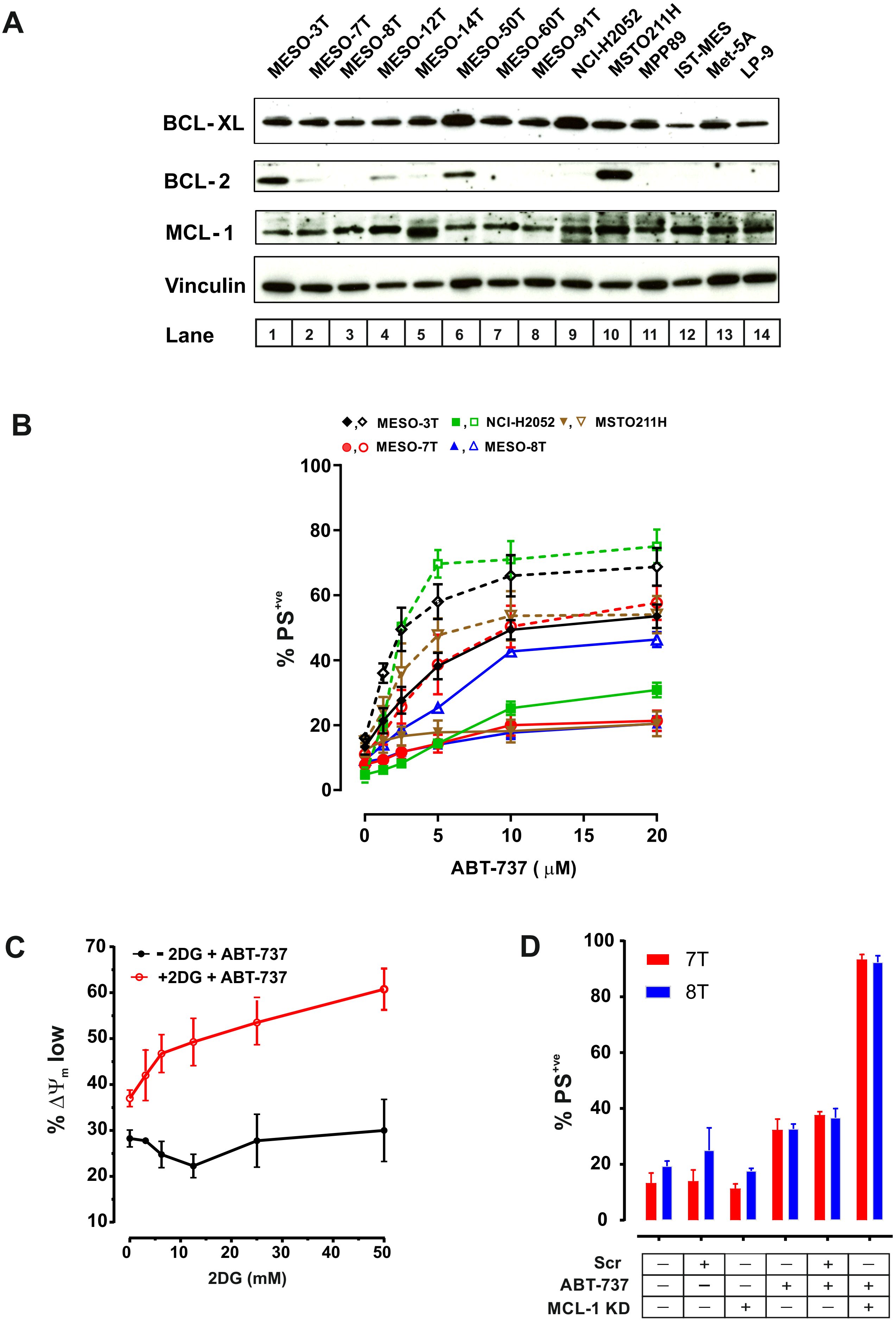

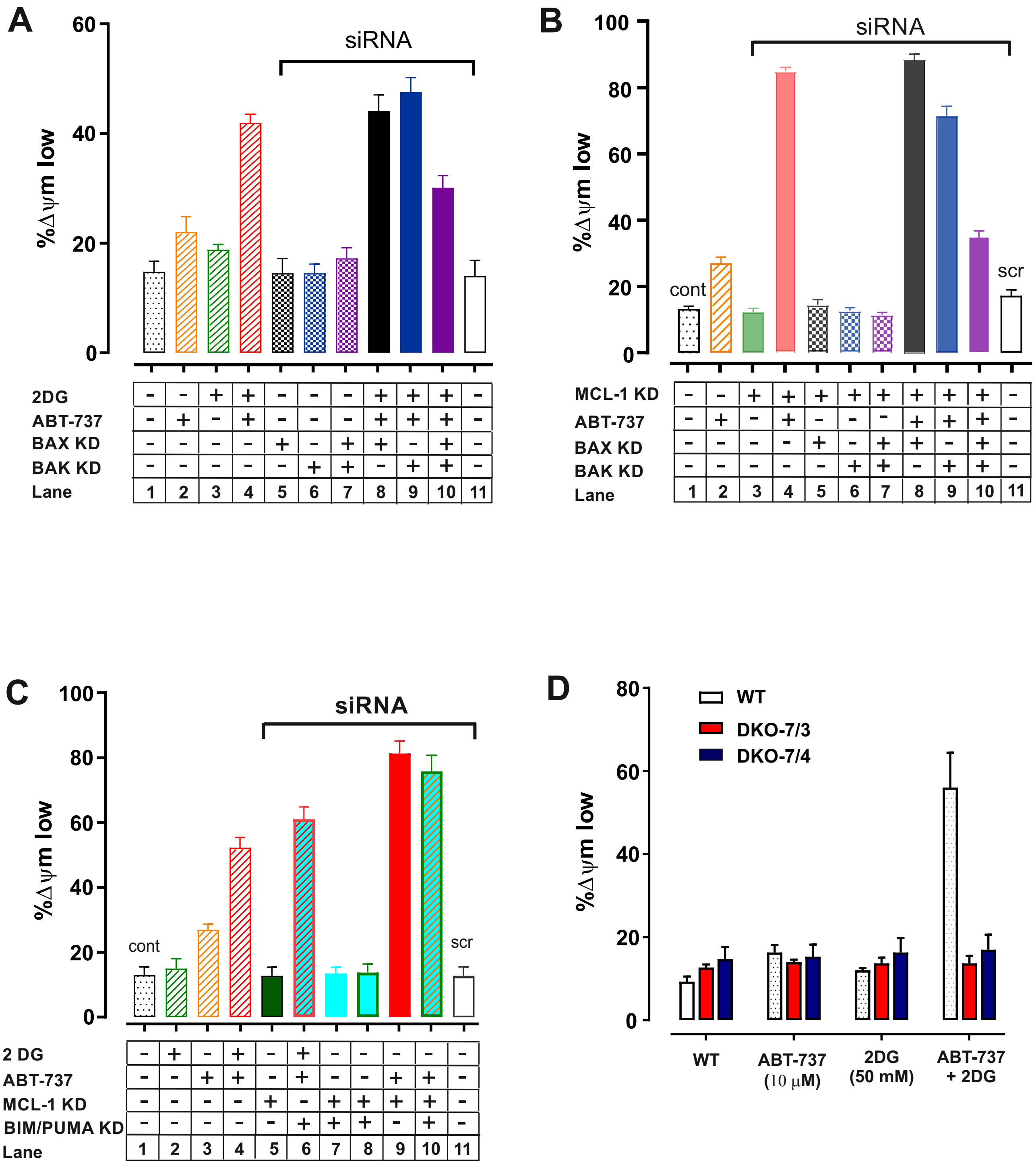

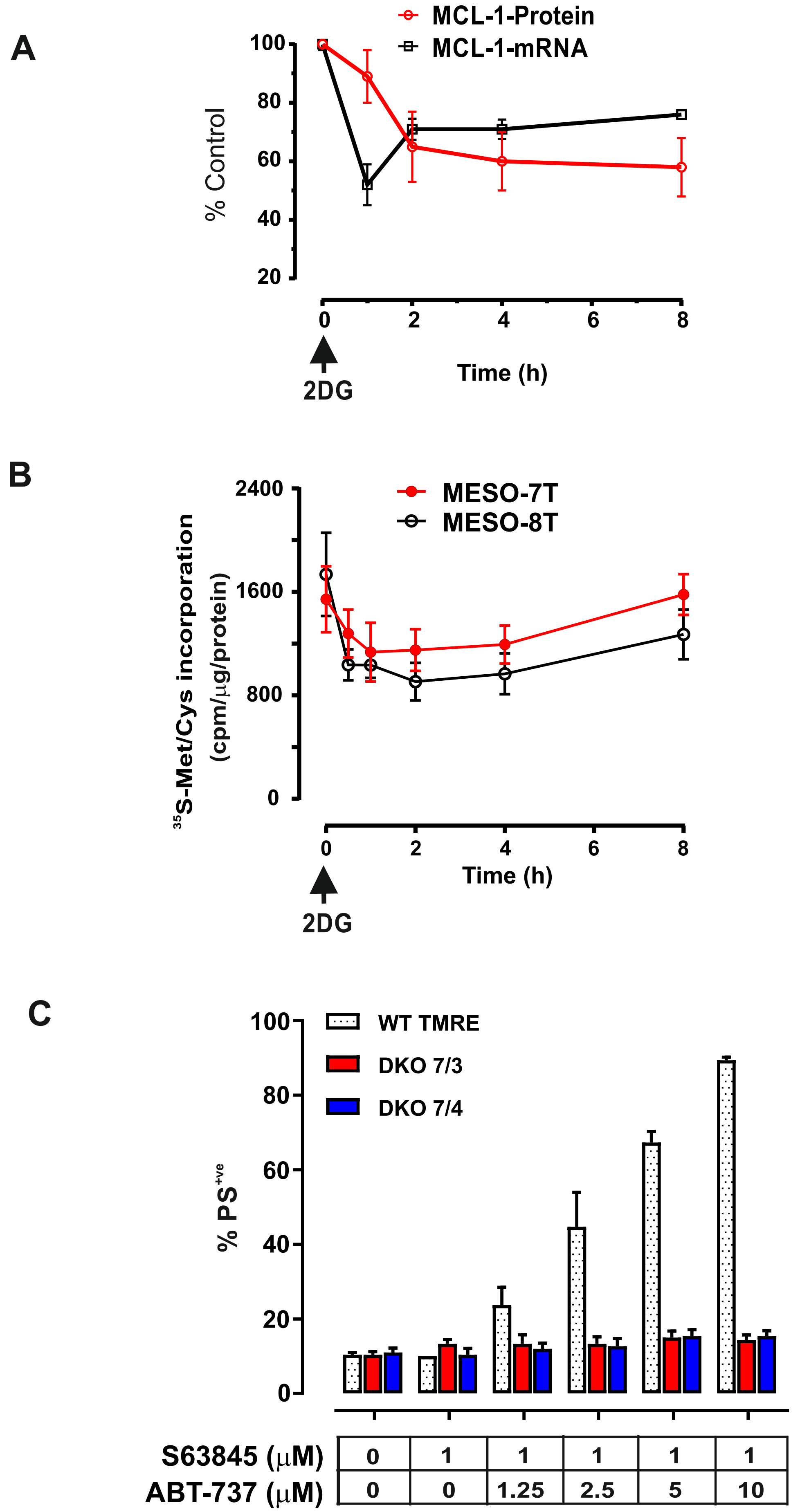

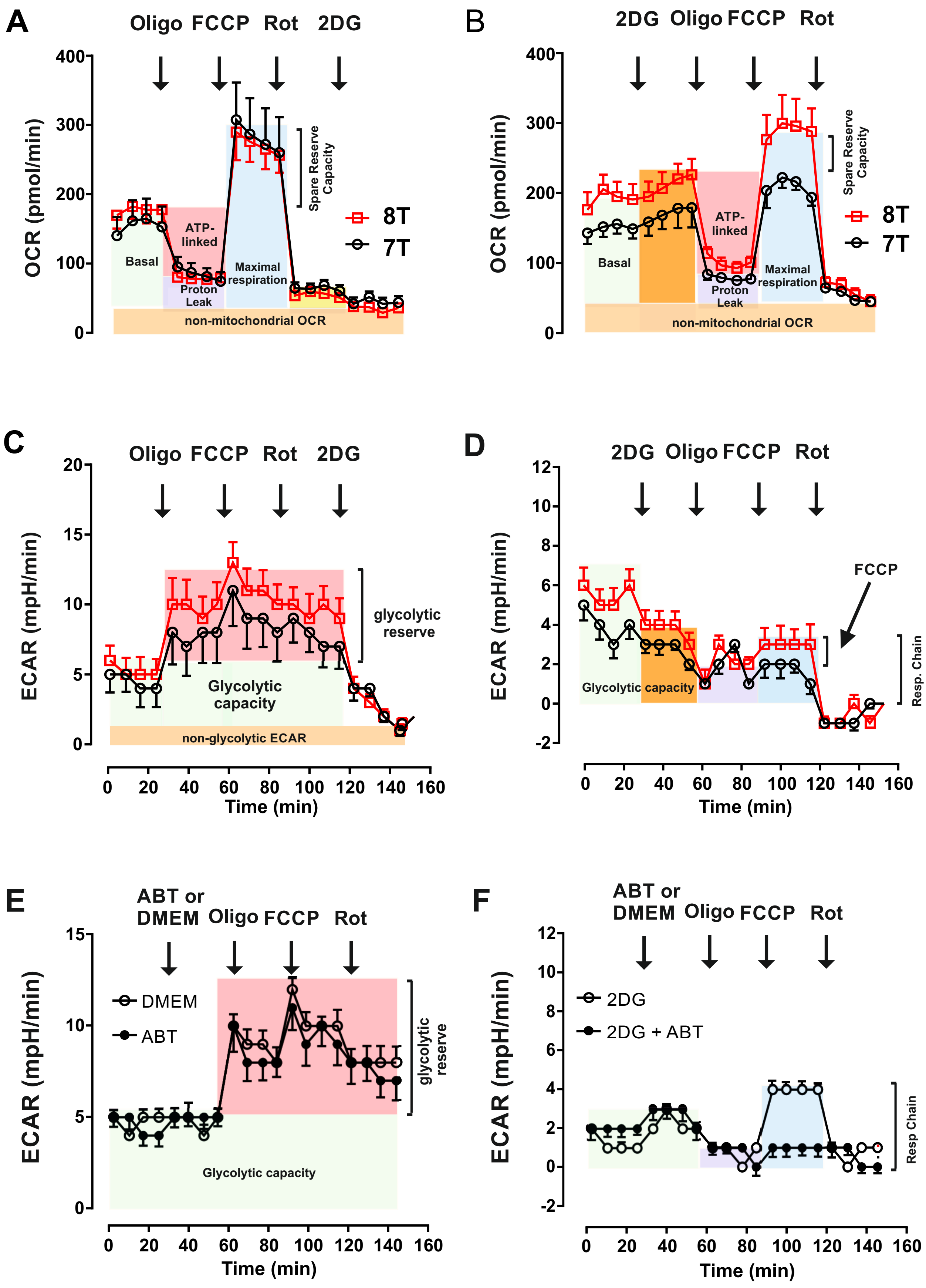

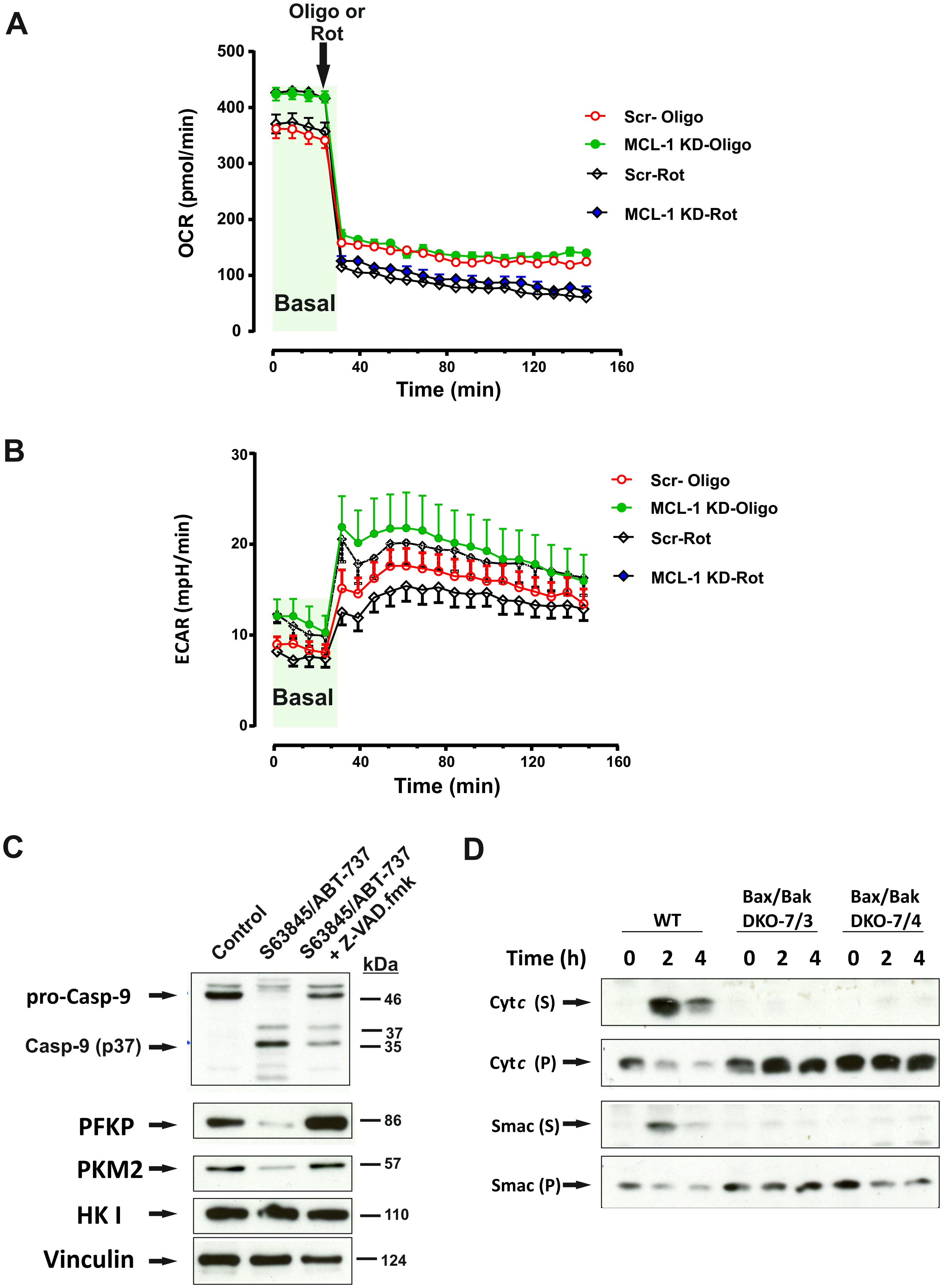

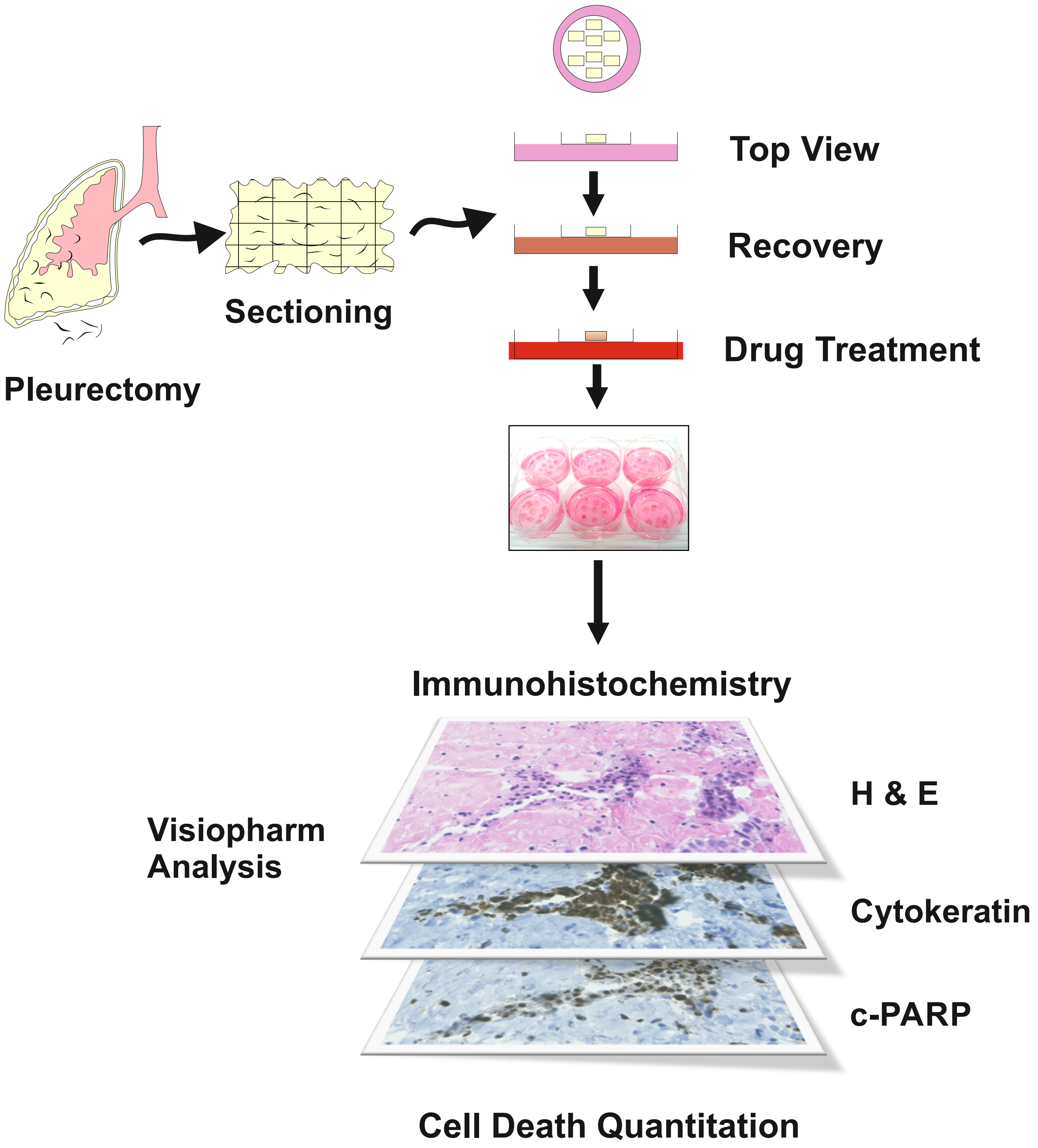
